# Genome-wide DNA methylation analysis of heavy cannabis exposure in a New Zealand longitudinal cohort

**DOI:** 10.1101/829598

**Authors:** Amy J. Osborne, John F. Pearson, Alexandra J. Noble, Neil J. Gemmell, L. John Horwood, Joseph M. Boden, Miles Benton, Donia P. Macartney-Coxson, Martin A. Kennedy

## Abstract

Cannabis use is of increasing public health interest globally. Here we examined the effect of cannabis use, with and without tobacco, on genome-wide DNA methylation in a longitudinal birth cohort (Christchurch Health and Development Study). We found the most differentially methylated sites in cannabis with tobacco users were in the *AHRR* and *F2RL3* genes, replicating previous studies on the effects of tobacco. Cannabis-only users had no evidence of differential methylation in these genes, or at any other loci at the epigenomewide significance level (P<10^−7^). However, there were 521 sites differentially methylated at P<0.001 which were enriched for genes involved in cardiomyopathy and neuronal signalling. Further, the most differentially methylated loci were associated with genes with reported roles in brain function (e.g. *TMEM190, MUC3L, CDC20* and *SP9*). We conclude that the effects of cannabis use on the mature human blood methylome differ from, and are less pronounced than, the effects of tobacco use, and that larger sample sizes are required to investigate this further.

## Introduction

Cannabis use is an important global public health issue, and a growing topic of controversy and debate ^1; 2^. It is the most widely used illicit psychoactive substance in the world ^3^, and the potential medicinal and therapeutic benefits of cannabis and its main active ingredients tetrahydrocannabinol (THC) and cannabidiol (CBD) are gaining interest^4–6^. There is strong evidence to suggest that the heavy and prolonged use of cannabis may be associated with increased risk of adverse outcomes in a number of areas, including mental health (psychosis ^7–9^, schizophrenia ^10; 11^, depression ^12; 13^), and illicit drug abuse ^14^.

Drug metabolism, drug response and drug addiction have known genetic components^15^, and multiple genome-wide association studies (GWAS) have identified genes and allelic variants that are likely contributors to substance use disorders^16; 17^. There are aspects of cannabis use disorder that are heritable ^18–21^, and several candidate loci for complex phenotypes such as lifetime cannabis use have recently been identified^3; 22^ that explain a proportion of the variance in cannabis use heritability. Complex phenotypes like these are influenced by multiple loci, each of which usually has a small individual effect size^23^, and such loci are frequently located in non-coding regions of the genome ^24; 25^, making their biological role difficult to elucidate.

Epigenetic mechanisms are involved in the interaction between the genome and environment; they respond to changes in environmental stimuli (such as diet, exercise, drugs), and act to alter chromatin structure and thus regulate gene expression^26^. Epigenetic modifications, such as DNA methylation, contribute to complex traits and diseases^27; 28^. Methylation of cytosine residues within CpG dinucleotides is an important mechanism of variation and regulation in the genome ^29–32^. Cytosine methylation, particularly in the promoter region of genes, is often associated with a decrease in transcription ^33^, and DNA methylation in the first intron and gene expression is correlated and conserved across tissues and vertebrate species^34^. Furthermore, modulation of methylation at CpG sites within the human genome can result in an epigenetic pattern that is specific to individual environmental exposures, and these may contribute to disease ^26; 35–37^. For example, environmental factors such as drugs, alcohol, stress, nutrition, bacterial infection, and exercise ^36; 38–41^ have been associated with methylation changes. A number of these methylation changes have been shown to endure and induce lasting biological changes^36^, whereas others are dynamic and transient. For example, alcohol consumption affects genome-wide methylation patterns in a severity-dependent manner^42^ and some of these changes revert upon abstinence from alcohol consumption ^43^. A similar observation is reported for former tobacco smokers with DNA methylation changes eventually reaching levels close to those who had never smoked tobacco after cessation ^44^. Thus, DNA methylation can be indicative of a particular environmental exposure, shed light on the dynamic interaction between the environment and the genome, and provide new insights in to the biological response.

Recreational drug use (an environmental stimuli) has been associated with adverse mental health outcomes particularly in youth ^45–49^, and epigenetics may play a role in mediating the biology involved. Therefore, we sought to determine whether regular cannabis users displayed differential cytosine methylation compared to non-cannabis users. Cannabis users in this study are participants from the Christchurch Health and Development Study (CHDS), a longitudinal study of a birth cohort of 1265 children born in 1977 in Christchurch, New Zealand. Users often consume cannabis in combination with tobacco. Unusually, the CHDS cohort contains a subset of cannabis users who have never consumed tobacco, thus enabling an investigation of the specific effects of cannabis consumption, in isolation, on DNA methylation in the human genome.

## Methods

### Cohort and study design

The Christchurch Health and Development Study includes individuals who have been studied on 24 occasions from birth to the age of 40 (n=987 at age 30, with blood collected at approximately age 28). In the early 1990s, research began into the initiation and consequences of cannabis use amongst CHDS participants; cannabis use was assessed prospectively over the period up to the collection of DNA^11–14; 48–54^. A subset of n=96 participants for whom a blood sample was available are included in the current study. Cases (regular cannabis users, n = 48) were matched with controls (n = 48) for sex (n=37 male, n=11 female each group, for additional information see Supplementary Table 1). Case participants were partitioned into two subsets: one that contained cannabis-only users (who had never consumed tobacco, “cannabis-only”, n = 24), and one that contained cannabis users who also consumed tobacco (“cannabis with tobacco”, n = 24) and were selected on the basis that they either met DSM-IV^55^ diagnostic criteria for cannabis dependence, or had reported using cannabis on a daily basis for a minimum of three years prior to age 28. Cannabis consumption was via smoking, for all participants. The median duration of regular use was 9 years (range 3-14 years). Control participants had never used cannabis or tobacco. Additionally, comprehensive SNP data was available for all participants^56^. All aspects of the study were approved by the Southern Health and Disability Ethics Committee, under application number CTB/04/11/234/AM10 “Collection of DNA in the Christchurch Health and Development Study”, and the CHDS ethics approval covering collection of cannabis use: “16/STH/188/AM03 The Christchurch Health and Development Study 40 Year Follow-up”.

### DNA extraction and methylation arrays

DNA was extracted from whole blood using the KingFisher Flex System (Thermo Scientific, Waltham, MA USA), as per the published protocols. DNA was quantified via NanoDrop™ (Thermo Scientific, Waltham, MA USA) and standardised to 100ng/μl. Equimolar amounts were shipped to the Australian Genomics Research Facility (AGRF, Melbourne, VIC, Australia) for analysis with the Infinium^®^ MethylationEPIC BeadChip (Illumina, San Diego, CA USA).

### Bioinformatics and Statistics

All analysis was carried out using R (Version 3.5.2 ^57^). Prior to normalisation, quality control was performed on the raw data. Firstly, sex chromosomes and 150 failed probes (detection P value > 0.01 in at least 50% of samples) were excluded from analysis. Furthermore, potentially problematic CpGs with adjacent SNVs, or that did not map to a unique location in the genome ^58^, were also excluded, leaving 700,296 CpG sites for further analysis. The raw data were then normalised with the NOOB procedure in the minfi package^59^ (Supplementary Figure 1). Normalisation was checked by visual inspection of intensity densities and the first two components from Multi-Dimensional Scaling of the 5000 most variable CpG sites (Supplementary Figures 2 and 3). The proportions of cell types (CD4+, CD8+ T cells, Natural Killer, B cells, Monocytes and Granulocytes) in each sample were estimated with the Flow.Sorted.Blood package^60^. Linear models were fitted to the methylated/unmethylated or M ratios using limma ^61^. Separate models were fitted for cannabis-only vs. controls, and cannabis plus tobacco users vs. controls. Both models contained covariates for sex (bivariate), socioeconomic status (three levels), batch (bivariate), population stratification (four principal components from 5000 most variable SNPs) and cell type (five continuous), β values were calculated, defined as the ratio of the methylated probe intensity (M) / the sum of the overall intensity of both the unmethylated probe (U) + methylated probe (M). P values were adjusted for multiple testing with the Benjamini and Hochberg method and assessed for genomic inflation with bacon^62^. Differentially methylated CpG sites were matched to the nearest neighbouring genes in Hg19 using GRanges^63^, and their official gene symbols were tested for enrichment in KEGG 2019 human pathways with EnrichR^64^.

## Results

### Data normalisation

Modelled effects showed no indication of genomic inflation with λ =1.04 for cannabis-only users (Supplementary Figure 4a) and λ = 0.855 for cannabis with tobacco users (Supplementary Figure 4b), versus controls. These were confirmed with bacon for cannabis-only (inflation = 0.98, bias = 0.044) and cannabis with tobacco users (inflation = 0.91, bias = 0.19). Inflation values less than 1 suggest that the results may be conservative.

Cannabis with tobacco users had a significantly lower estimated proportion of natural killer cells than controls (1.8%, 0.4% - 3.2%, P<0.014) with no other proportions differing significantly. After adjusting for multiple comparisons this was not significant (P=0.08) however we note that it is consistent with other findings that NK-cells are suppressed in the plasma of tobacco smokers^65; 66^.

### Differential methylation

The most differentially methylated CpG sites for cannabis users relative to controls differed in the absence (Table 1) and presence (Table 2) of tobacco smoking. Five individual CpG sites were significantly differentially methylated (P adjusted <0.008) between users and controls when cannabis with tobacco was used (Table 2 and Figure 1). The top CpG sites in the *AHRR, ALPG* and *F2RL3* genes (Table 2) are consistent with previous studies on tobacco use without cannabis (e.g. ^44;67–69^), and cg17739917 is in the same CpG-island as other CpGs previously shown to be hypomethylated in response to tobacco^70^. Cannabis-only users showed no CpG sites differentially methylated after correction for multiple testing (Table 1 and Figure 2), however the most differentially methylated site was hypermethylation of cg12803068 in the gene *MYO1G*, which has been reported to be hypermethylated in response to tobacco use^67^.

**Table 1 –.** Top 15 differentially methylated CpG sites in cannabis-only users vs controls.. Beta values with P values, nominal and adjusted by the Benjamini and Hochberg method. Locations are relative to hg19 with gene names for overlapping genes or nearest 5’ gene with distance to the 5’ end shown.

| CpG | Gene | Location | Distance | Cannabis | Control | Difference | P value | P value |
| --- | --- | --- | --- | --- | --- | --- | --- | --- |
| | | | | $\beta_U$ | $\beta_C$ | $\beta_U - \beta_C$ | *Nominal* | *Adjusted* |
| cg12803068 | MYO1G | intron |  | 0.8 | 0.71 | 0.1 | 6.30E-07 | 0.4 |
| cg02234936 | ARHGEF1 | intron |  | 0.14 | 0.13 | 0.01 | 1.10E-06 | 0.4 |
| cg01695406 | TMEM190 | intron |  | 0.82 | 0.77 | 0.05 | 3.00E-06 | 0.6 |
| cg24875484 | MUCL3 | intron |  | 0.1 | 0.09 | 0.01 | 3.90E-06 | 0.6 |
| cg05009104 | MYO1G | intron |  | 0.79 | 0.74 | 0.05 | 5.90E-06 | 0.6 |
| cg00470351 | CDC20 | exon |  | 0.4 | 0.38 | 0.02 | 6.10E-06 | 0.6 |
| cg24060040 | DUS3L | upstream | 11,018 | 0.11 | 0.08 | 0.03 | 6.30E-06 | 0.6 |
| cg12322720 | FOXB1 | downstream | 150,921 | 0.58 | 0.52 | 0.06 | 8.90E-06 | 0.7 |
| cg16746471 | KIAA1324L | promoter | 374 | 0.1 | 0.08 | 0.02 | 1.10E-05 | 0.7 |
| cg04180046 | MYO1G | intron |  | 0.56 | 0.52 | 0.04 | 1.20E-05 | 0.7 |
| cg06955687 | DDX25 | downstream | 28,769 | 0.74 | 0.7 | 0.04 | 1.20E-05 | 0.7 |
| ch.22.70704 | TNRC6B | downstream | 159,737 | 0.06 | 0.04 | 0.01 | 1.30E-05 | 0.7 |
| cg09344183 | SP9 | downstream | 5,964 | 0.06 | 0.05 | 0.01 | 1.40E-05 | 0.7 |
| cg06693983 | TMEM190 | exon |  | 0.84 | 0.76 | 0.08 | 1.40E-05 | 0.7 |
| cg26069230 | ADAP2 | exon |  | 0.16 | 0.14 | 0.01 | 1.50E-05 | 0.7 |

**Table 2.**
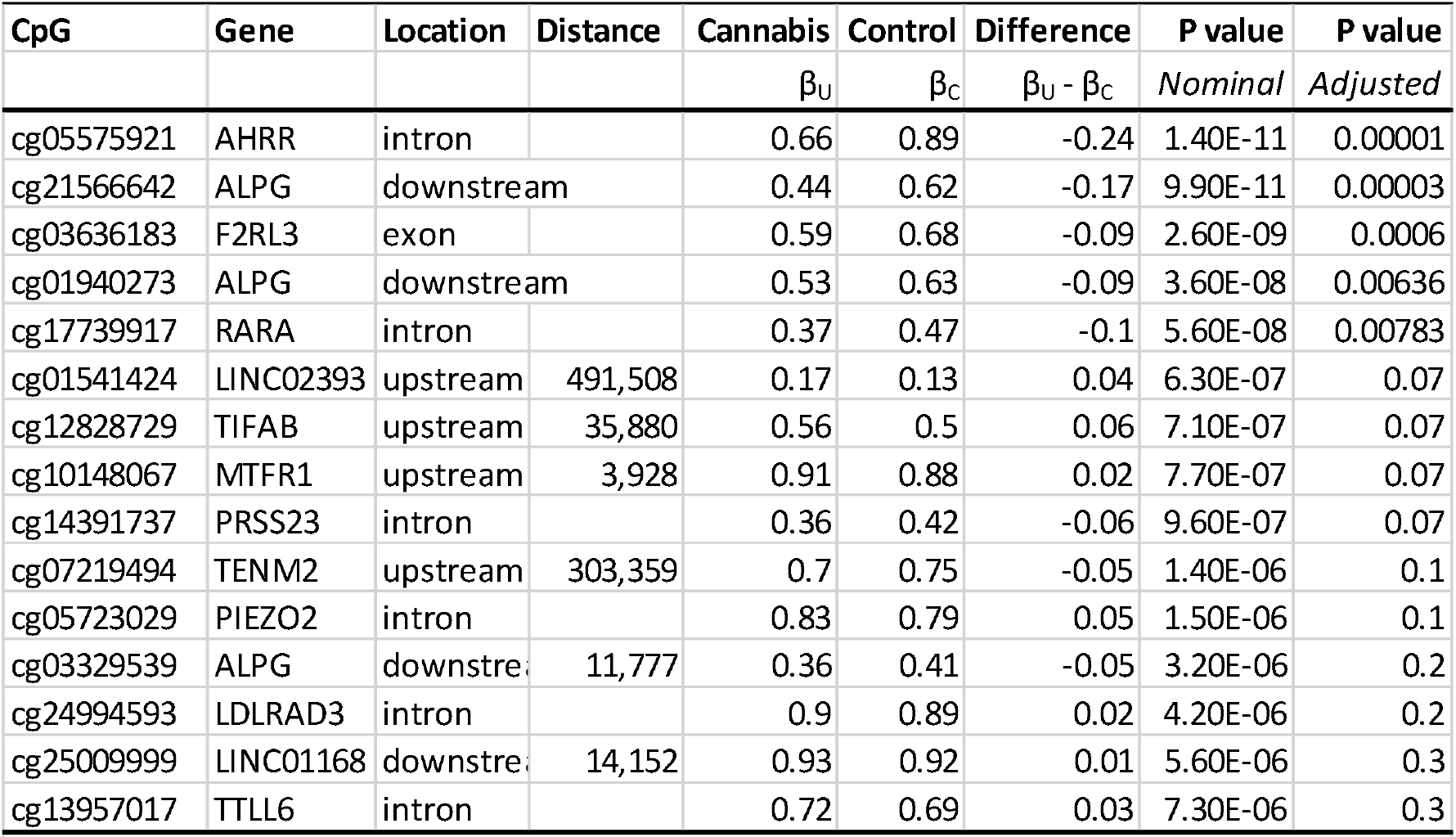
Top 15 differentially methylated CpG sites in cannabis with tobacco users vs controls. Beta values with P values, nominal and adjusted by the Benjamini and Hochberg method. Locations are relative to hg19 with gene names for overlapping genes or nearest 5’ gene with distance to the 5’ end shown.

**Figure 1 –.**
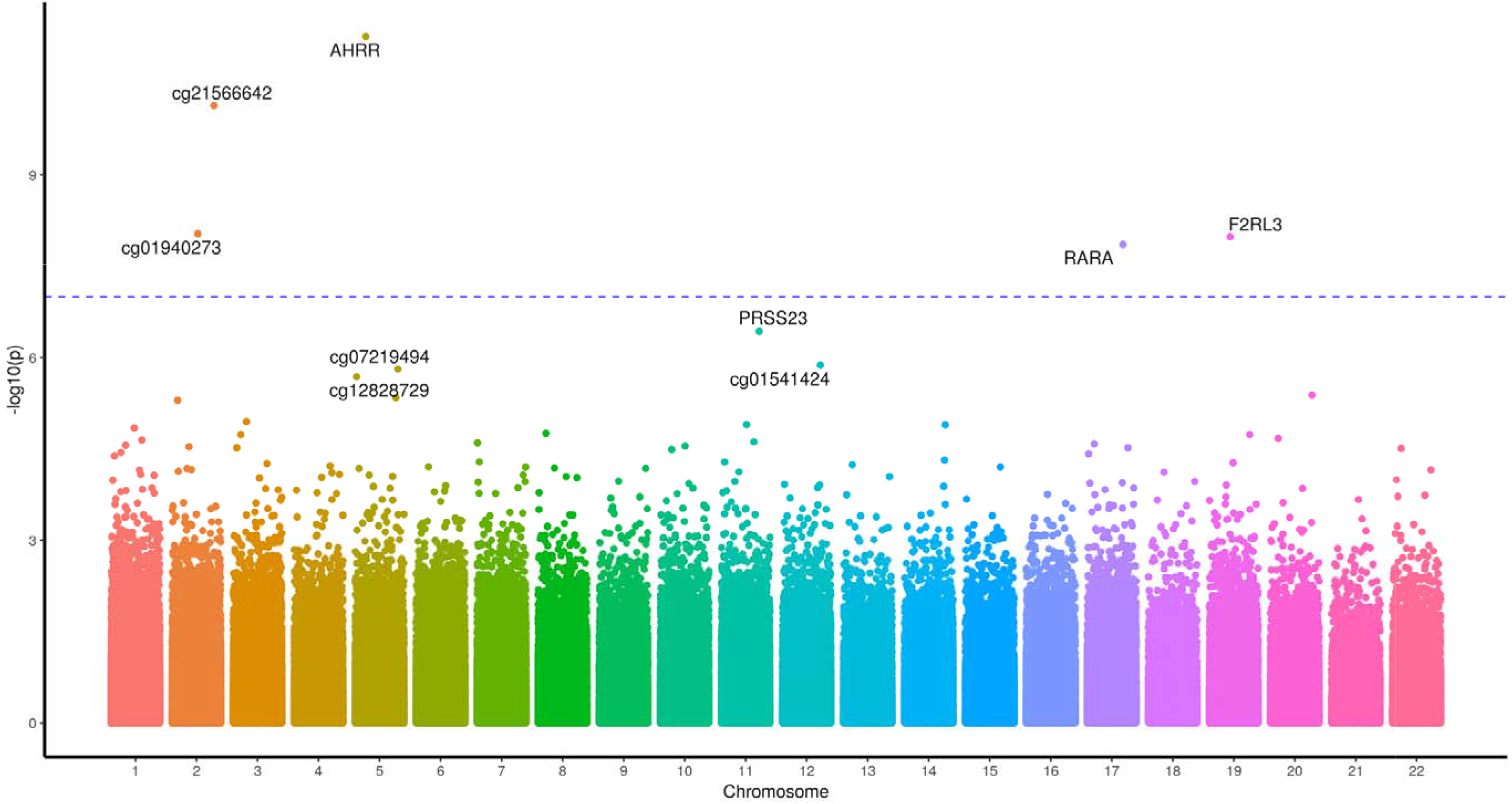
A Manhattan plot of the genome-wide CpG sites found in the cannabis with tobacco analysis. The Y axis presents −log10(p) values with the most significant methylated sites labelled with the gene the CpG site resides in.

**Figure 2 -.**
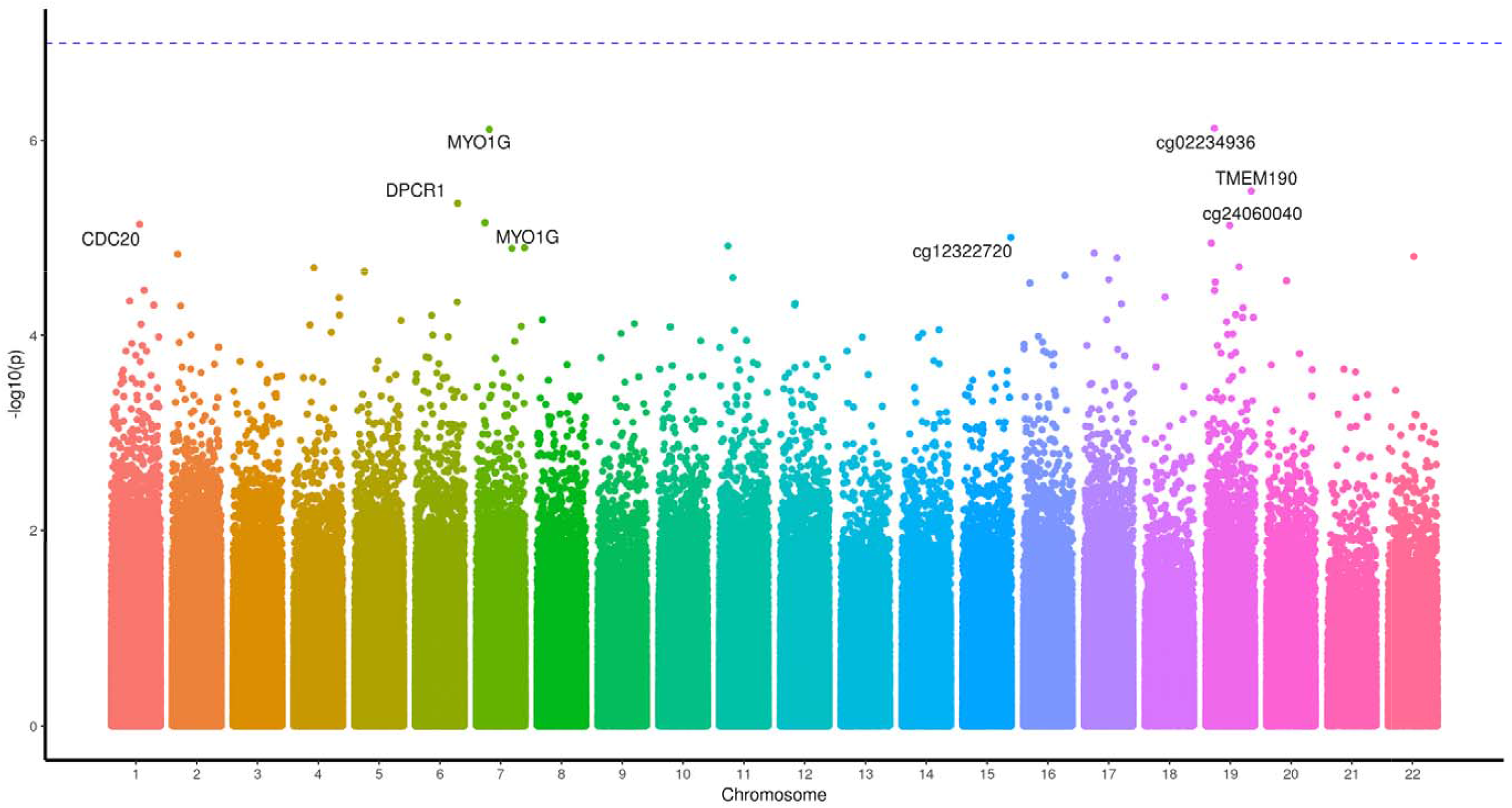
A Manhattan plot of the genome wide CpG sites found in the cannabis-only analysis. The Y axis presents −log10(p) values with the most nominally significant different methylated sites labelled with the gene the CpG site resides in.

To describe the data we chose a nominal P value of 0.001, and observed that both cannabis-only and cannabis with tobacco users showed relatively higher rates of hypermethylation than hypomethylation compared to controls and that the distribution of these CpG sites was similar with respect to annotated genomic features (Table 3). Four CpG sites overlapped between the cannabis-only and cannabis with tobacco users analyses; two were hypermethylated; cg02514528, in the promoter of *MARC2*, and cg27405731 in *CUX1*, and one, cg26542660 in the promoter of *CEP135*, was hypomethylated in comparison to controls. The second most differentially methylated site (ranked by P value) in cannabis-only users was cg02234936 which maps to *ARHGEF1*; this was hypermethylated in the cannabis with tobacco users.

**Table 3.** Summary of CpG sites from cannabis-only and cannabis with tobacco users vs. nonusers. Counts of significant sites at P= 0.001 and at a Benjamini and Hochberg adjusted P < 0.05. ‘Both’ indicates the number of CpG sites of each type that are present and shared across both analyses.

|  | Cannabis only |  | Tobacco + Cannabis |  | Both |
| --- | --- | --- | --- | --- | --- |
| **Differentially Methylated loci (FWER = 0.05)** | 0 |  | 6 |  |  |
| **Differentially Methylated loci (P<0.001)** |  |  |  |  |  |
| Total | 521 |  | 533 |  |  |
| Hypermethylated | 420 | 80.6% | 403 | 75.6% | 2 |
| Hypomethylated | 101 | 19.4% | 130 | 24.4% | 1 |
| Hyper (Cannabis) Hypo (Cannabis + Tobacco) |  |  |  |  | 1 |
| **Location** |  |  |  |  |  |
| Intron | 216 | 41.5% | 264 | 49.5% |  |
| Exon | 97 | 18.6% | 65 | 12.2% |  |
| Exon Boundary | 0 |  | 0 |  |  |
| Promoter | 89 | 17.1% | 60 | 11.3% |  |
| 3' UTR | 3 | 0.6% | 1 | 0.2% |  |
| 5' UTR | 0 |  | 0 |  |  |
| 3' (downstream) | 62 | 11.9% | 76 | 14.3% |  |
| 5' (upstream) | 54 | 10.4% | 67 | 12.6% |  |

### Pathway enrichment analyses

We then took the genes containing differentially methylated CpG sites at P<0.001 for the cannabis-only group that were within genes (that is, not up or downstream in Table 3) and compared them with human KEGG pathways using Enrichr. The hypermethylated CpG sites (n = 420) showed enrichment in the arrhythmogenic right ventricular cardiomyopathy pathway at an adjusted P = 0.03 and enrichment in the glutamatergic synapse and long term potentiation pathway at an adjusted P=0.05 (Figure 3). Enrichment analysis of hypomethylated loci (n = 101) in cannabis-only users did not identify any KEGG pathways at or near adjusted significance (P=0.05, Figure 4).

**Figure 3 –.**
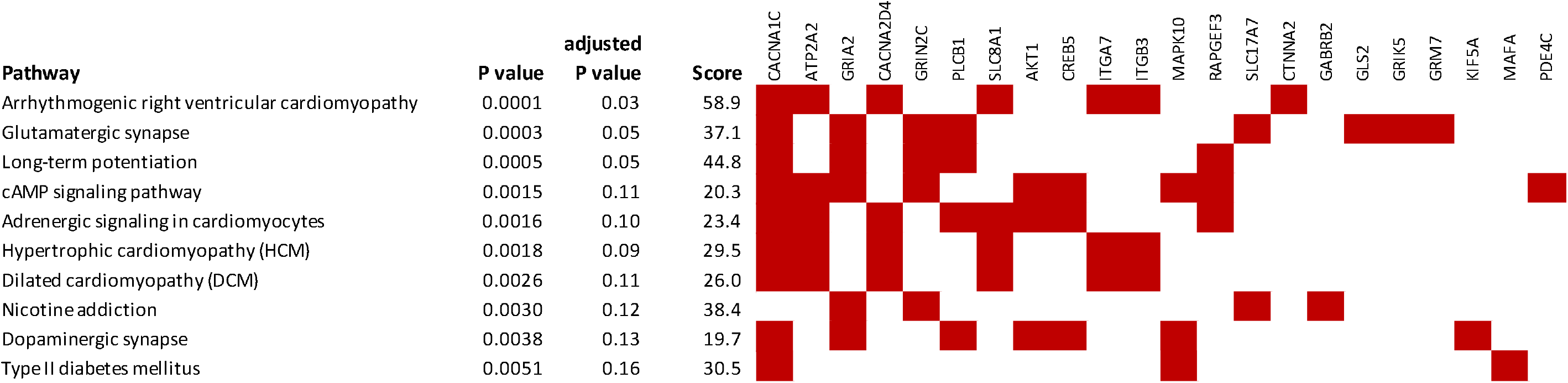
Genetic networks enriched within the hypermethylated CpG sites identified in the cannabis-only analysis. Pathways from KEGG 2019. Genes shown by filled cells are hypermethylated in cannabis-only users and included in named pathway.

**Figure 4 –.**
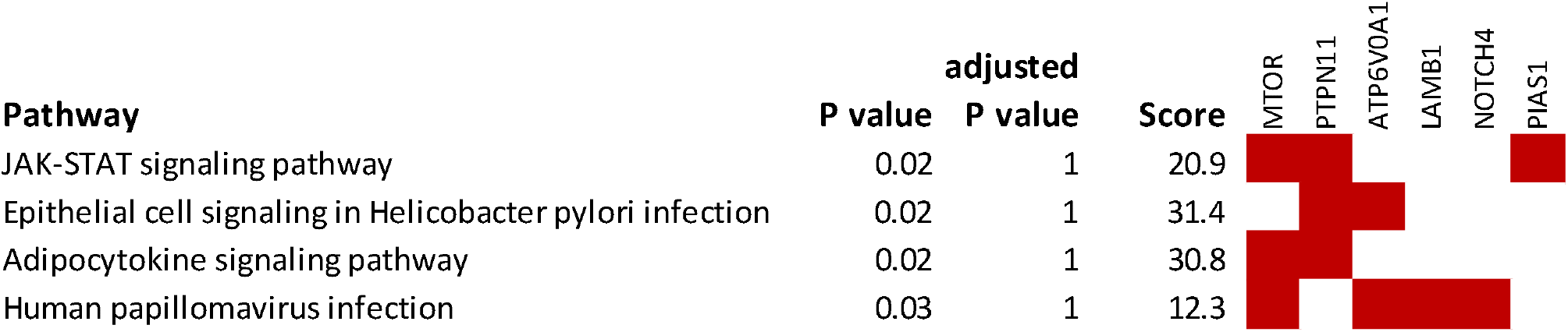
Genetic networks enriched within the hypomethylated CpG sites identified in the cannabis-only users. Pathways from KEGG 2019. Genes shown by filled cells are hypomethylated in cannabis-only users and included in named pathway.

## Discussion

Many countries have recently adopted, or are considering, lenient polices regarding the personal use of cannabis ^71–73^. This approach is supported by the evidence that the prohibition of cannabis can be harmful^53^. Further, the therapeutic benefits of cannabis are gaining traction, most recently as an opioid replacement therapy^74^. However, previous studies, including analyses of the CHDS cohort, have reported an association between cannabis use and poor health outcomes, particularly in youth ^75; 76^. Epigenetic mechanisms, including DNA methylation, provide the interface between the environment (e.g. cannabis exposure) and genome. Therefore, we investigated whether changes in an epigenetic mark, DNA methylation, were altered in cannabis users, versus controls, a comparison made possible by the deep phenotyping of the CHDS cohort with respect to cannabis use, and the fact that the widespread practice of mulling or mixing cannabis with tobacco, is not common in New Zealand.

Consistent with previous reports of tobacco exposure, we observed greatest differential methylation in cannabis with tobacco users in the *AHRR* and *F2RL3* genes ^44;67–69^. These changes, however, were not apparent in the cannabis-only data. Only two nominally significantly differentially methylated (P<0.05) CpG sites were observed in both the cannabis-only and cannabis with tobacco analyses. This suggests that tobacco may have a more pronounced effect on DNA methylation and/or dominates any effects of cannabis on the human blood methylome, and that caution should be taken when interpreting similar cannabis exposure studies which do not, or cannot, exclude tobacco smokers. Interestingly, the two nominally significant CpG sites (P<0.05) that overlap between the cannabis-only and the cannabis with tobacco data are located within the *MARC2* and *CUX1* genes, which both have reported roles in brain function; a SNP in *MARC2* has been provisionally associated with the biological response to antipsychotic therapy in schizophrenia patients^77^, and the *CUX1* gene has an established role in neural development^78^.

Cannabis affects the brain, leading to perceptual alterations, euphoria and relaxation^18^, and prolonged use is associated with mood disorders, including adult psychosis ^7; 8; 49; 79; 80^, mania ^13^, and depression ^12^. We did not detect significantly differentially methylated loci associated with exclusive cannabis use at the epigenome-wide level. However, an assessment of those top loci reaching nominal significance (P<0.05) identified CpG sites within genes involved in brain function and mood disorders, including *MUC3L*^81; 82^, *CDC20* ^83^, *DUS3L* ^84^ *TMEM190* ^85^, *FOXB1* ^86–88^, *KIAA1324L/GRM3* ^82; 89–94^, *DDX25* ^81; 95; 96^ *TNRC6B* ^97; 98^ and *SP9^99^*.

Pathway enrichment revealed that hypermethylation in cannabis-only users was overrepresented in genes associated with cardiomyopathies and neural signalling. This is consistent with the literature which raises clinical concerns around cardiac complications potentially associated with cannabis use ^100–103^. The enrichment of genes associated with neural signalling pathways is also consistent with the literature, including previous analyses of the CHDS cohort, which report associations between cannabis exposure and brain related biology such as mood disorders^7; 12; 48; 49; 51–54; 104; 105^. Our study was limited by sample size, achieving approximately 10% power at P=10^−7^ to detect the largest standardized effect size found. However, while we have not implicated any gene at the genome-wide significance level with respect to differential methylation associated with cannabis-only exposure, our data is strongly suggestive of a role for DNA methylation in the biological response to cannabis, a possibility which definitely warrants further investigations in larger cohorts.

In summary, while tobacco use has declined on the back of state-sponsored cessation programs ^106^, rates of cannabis use remain high in New Zealand and globally, and might be predicted to increase further with the decriminalisation or legalisation of cannabis use for therapeutic and/or recreational purposes ^107^. Therefore, analysis of the potential effects of cannabis (an environmental stimuli) on DNA methylation, an epigenetic mechanism, is timely. Our data is strongly suggestive of a role for DNA methylation in the biological response to cannabis, significantly contributes to the growing literature studying the biological effects of heavy cannabis use, and highlights areas of further analysis in particular with respect to the epigenome.

## Acknowledgements

Allison Miller for technical assistance. Funding: CHDS, University of Otago Division of Health Sciences Collaborative Postdoctoral Fellowship to AO, University of Otago Research Grant to MK, The Carney Centre for Pharmacogenomics. CHDS funded by the Health Research Council of New Zealand (Programme Grant 16/600) and the Canterbury Medical Research Foundation.

