## Supplementary for "Genome-wide DNA methylation analysis of heavy cannabis exposure in a New Zealand longitudinal cohort"

|  |  | **Cases** | **Controls** |
| --- | --- | --- | --- |
| Sex | Male | 37 | 37 |
|  | Female | 11 | 11 |
| Ethnicity | European | 35 | 45 |
|  | Other | 13 | 3 |
| Socioeconomic status | Professional/managerial | 6 | 6 |
|  | Clerical/technical/skilled | 21 | 21 |
|  | Semi-skilled/unskilled | 21 | 21 |

**Supplementary Table 1.** Christchurch Health and Development Study (CHDS) participants selected for EPIC arrays. Cases and controls were matched as closely as possible by sex, ethnicity and parental socioeconomic status/occupation. Cases are comprised of regular cannabis users, half of whom have never used tobacco. Controls are comprised of individuals with no exposure to cannabis or tobacco. ‘Other’ ethnicity is a combination of Māori and Pacific Island participants.

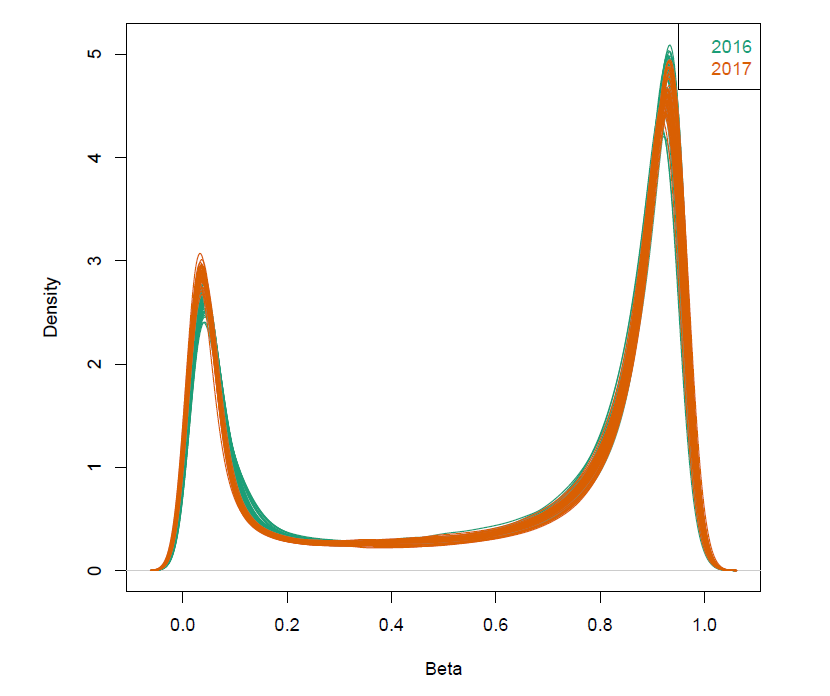

**Supplementary Figure 1. Preprocessing using noob normalisation.** The distribution of individuals as displayed as lines were plotted based on beta density post normalisation using noob. Data coloured based on year the measurements took place; 2016 (green) or 2017 (orange).

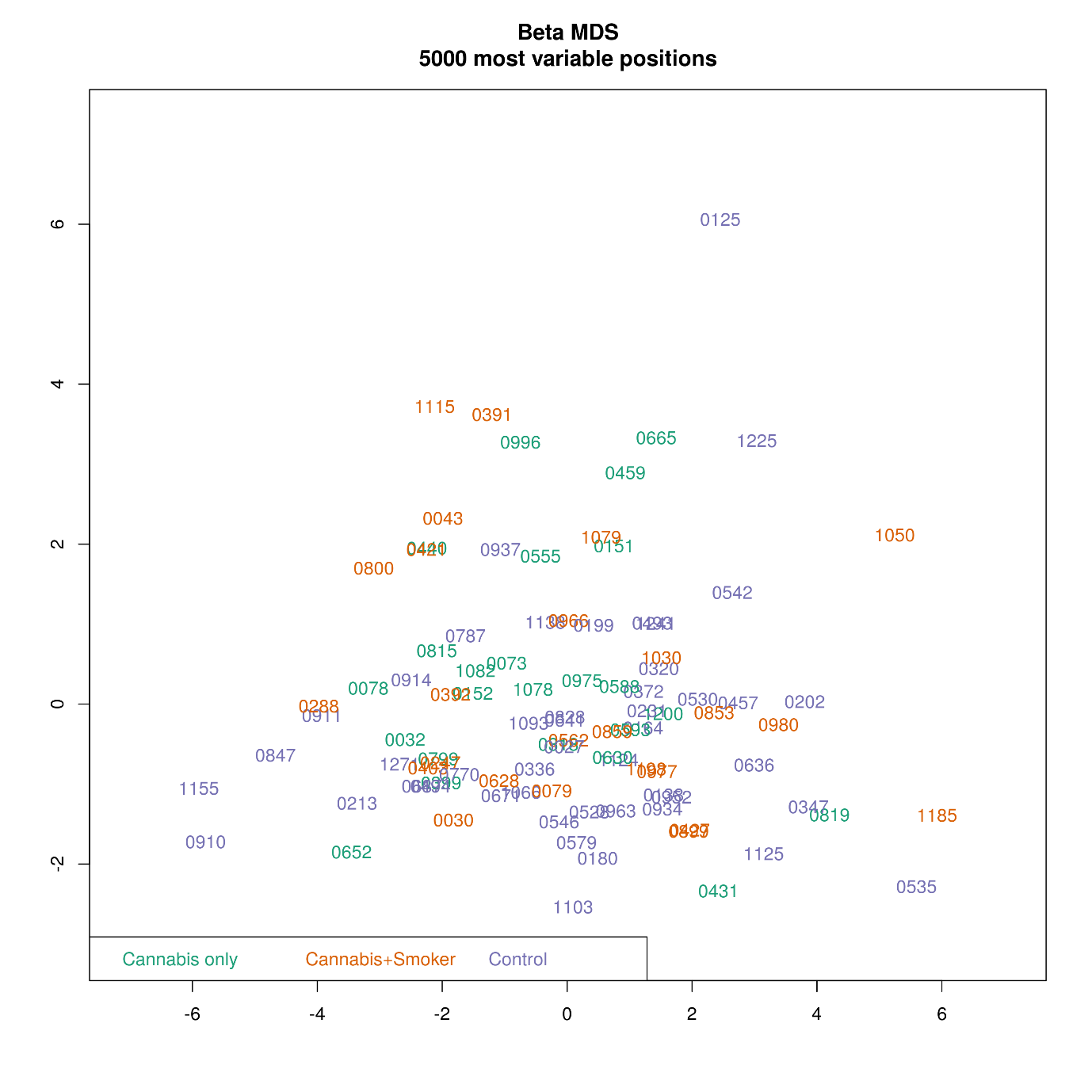

**Supplementary Figure 2. A multidimensional plot displaying the 5000 most variable positions post normalisation using noob.** Individuals are grouped in colour by status: cannabis-only users (green), cannabis with tobacco (orange) or control (purple).

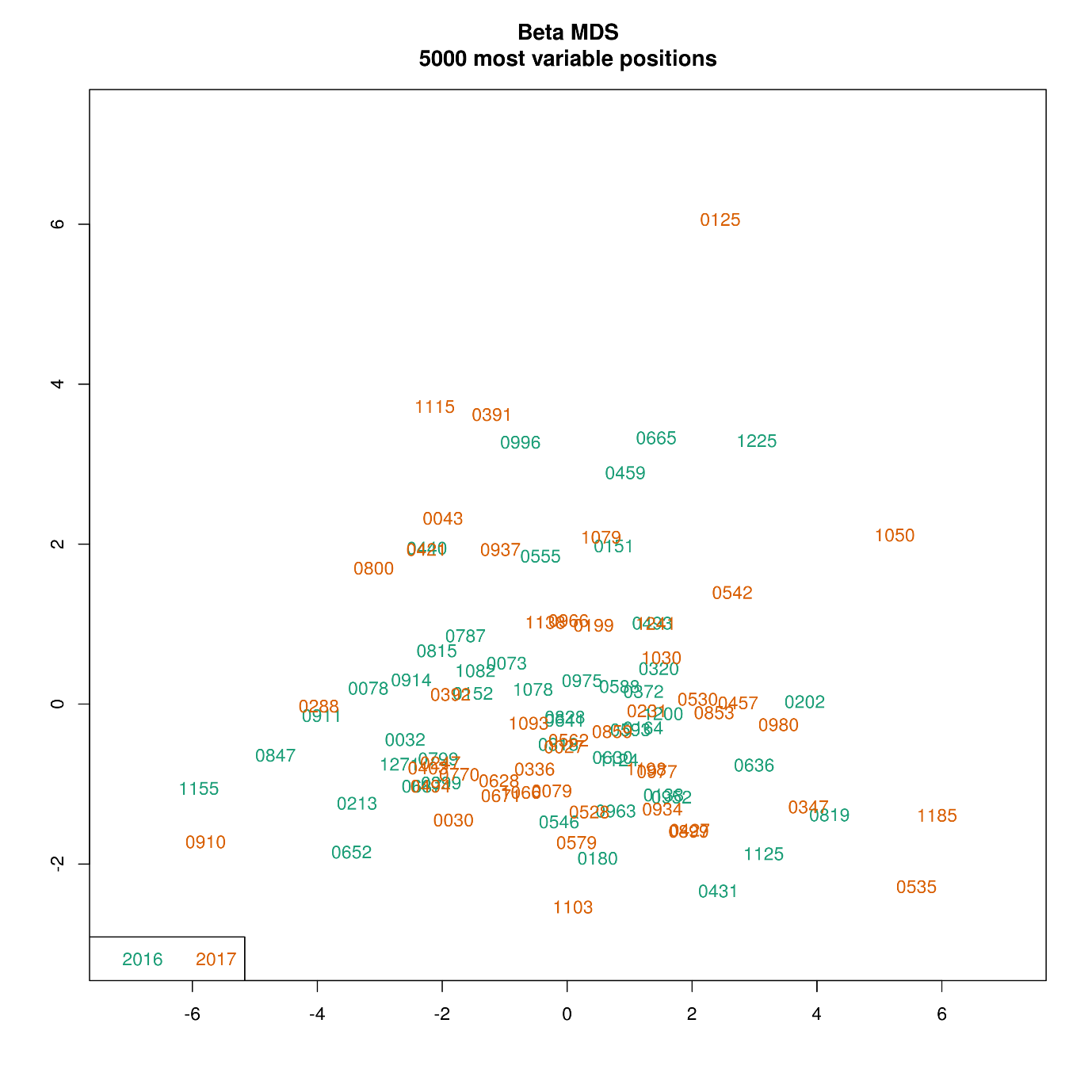

**Supplementary Figure 3. A multidimensional plot displaying the 5000 most variable positions post normalisation using noob.** Individuals are coloured on year the measurements were collected.

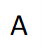

B

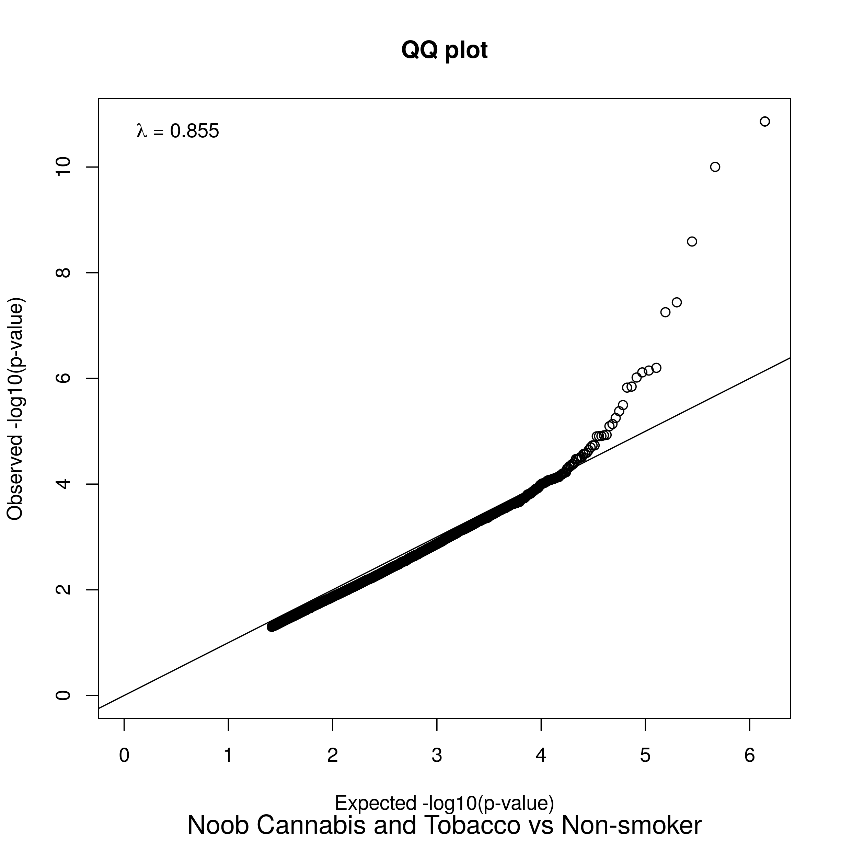

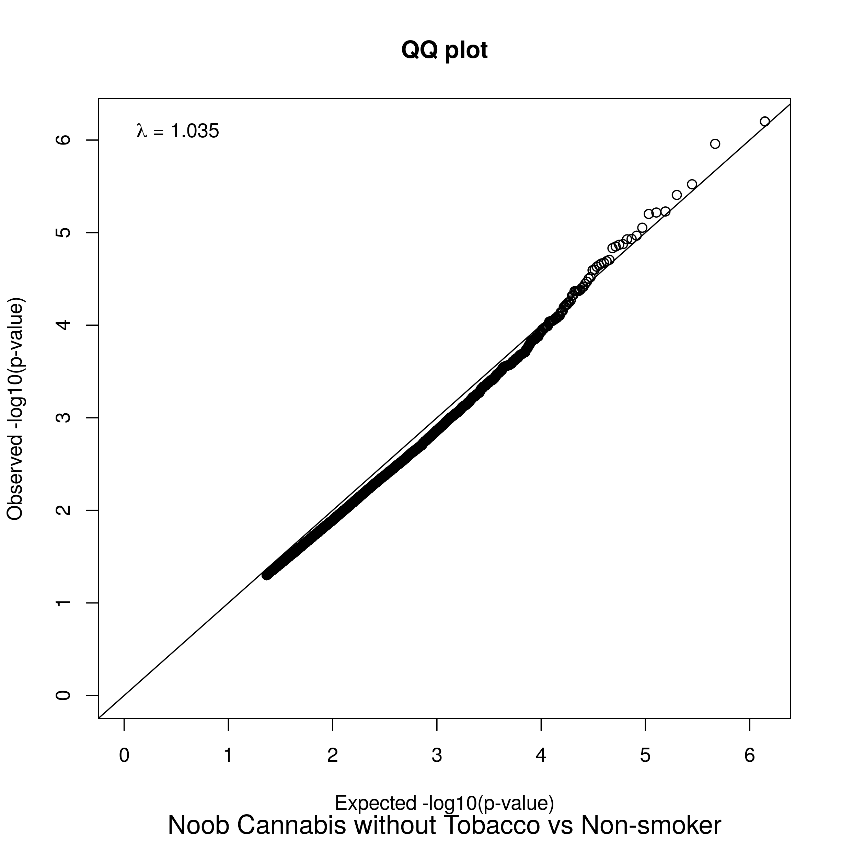

**Supplementary Figure 4. Quantile- quantile plots.** Quantile plots were used to assess for overfitting of models. A) Cannabis-only vs control samples B) Cannabis with tobacco vs control. Each dot displays the expected –log10 (p-values) under the model.
